# Metabolic and spatio-taxonomic response of uncultivated seafloor bacteria following the Deepwater Horizon oil spill

**DOI:** 10.1101/084707

**Authors:** KM Handley, YM Piceno, P Hu, LM Tom, OU Mason, GL Andersen, JK Jansson, JA Gilbert

**Author notes:** Correspondence should be addressed to Jack A. Gilbert, Ph.D. Department of Surgery, University of Chicago, 5842 South Maryland Avenue, Chicago, IL, 60637, USA; and Kim M. Handley, School of Biological Sciences, University of Auckland, Auckland, New Zealand.

## Abstract

The release of 700 million liters of oil into the Gulf of Mexico over a few months in 2010 produced dramatic changes in the microbial ecology of the water and sediment. Previous 4 studies have examined the phylogeny and function of these changes, but until now a 5 fundamental examination of the extant hydrocarbon metabolisms that supported these 6 changes had not been performed. Here, we reconstructed the genomes of 57 widespread 7 uncultivated bacteria from post spill sediments, and recovered their gene expression 8 pattern across the seafloor. These genomes comprised a common collection of bacteria 9 that were highly enriched in heavily affected sediments around the wellhead. While rare in distal sediments, some members were still detectable at sites up to 60 km away. Many of these genomes exhibited phylogenetic clustering indicative of common trait selection by the environment, and within half we identified 264 genes associated with hydrocarbon degradation. Observed alkane degradation ability was near ubiquitous among candidate hydrocarbon degraders, while just 3 harbored elaborate gene inventories for the degradation of alkanes and (poly)aromatic hydrocarbons. Differential gene expression profiles revealed a spill-promoted microbial sulfur cycle alongside gene up-regulation associated with polyaromatic hydrocarbon degradation. Gene expression associated with alkane degradation was widespread, although active alkane degrader identities changed along the pollution gradient. The resulting analysis suggests a broad metabolic capacity to respond to oil exists across a large array of usually rare bacteria.

---

Marine oil spills are frequent occurrences that can have a severe impact on environmental health and dependent economies^1^. In US marine environments alone, hundreds of oil spills occur annually, releasing millions of liters of oil per year on average^2^. While spills have typically occurred at shallow water depths, an expansion of drilling into the deep-sea led to the 2010 Deepwater Horizon (DWH) accident in the Gulf of Mexico^3^. DWH released over 700 million liters of oil from the Macondo MC252 wellhead at 1500 m depth^4^. This yielded a vast sea surface oil slick^5^, and an expansive plume of hydrocarbons at a water column depth of ~1100 m^6^. The leak also polluted deep-sea sediments up to tens of kilometers distance, due to direct contamination and flocculent fallout from plumes^7^. Pollution of seafloor sediments persisted at least 3 months post spill^8^, likely supported by ongoing inputs from sinking hydrocarbon-bearing particles^9^. Post spill contamination of sediment was greater than in the water column and was greatest within 3 km of MC252, where total polyaromatic hydrocarbon (PAH) concentrations remained above the Environmental Protection Agency's Aquatic Life benchmark^8,10^.

Natural bacterial communities are important agents for breaking down the complex mixtures of hydrocarbons in leaked oil^11,12^. Extensive degradation of petroleum hydrocarbons released during the DWH spill has been largely attributed to microbial activity^13^. Gammaproteobacteria – some clearly related to psychrotolerant or psychrophilic bacteria – dominated the deep-sea response^10,14,15^, contrasting with higher alphaproteobacterial ratios in shallow environments^16^. Although the buoyant plume received far greater attention than the underlying seafloor sediments^13^, post-spill sediments were observed to share some key taxa with the plume; a bacterium associated with the hydrocarbonoclastic genus *Colwellia* and an abundant undescribed gammaproteobacterium^10^. Genes or transcripts associated with hydrocarbon degradation were enriched in both the plume and polluted sediments^10,17,18^, and some were linked to single cell genomes of two plume alkane-degrading *Colwellia* and Oceanospirillales bacteria^18,19^. However, the communitywide and organism-specific metabolic response to the spill has not been directly explored leaving much of the hydrocarbon degrading potential of numerous uncultivated oil-responsive bacteria enigmatic, particularly on the seafloor.

Here we used 13 metagenomes^10^ and 10 metatranscriptomes to link communitywide hydrocarbon degradation strategies with microbial taxonomy in the top one centimeter of deep-sea sediment, collected 3 months after the DWH wellhead was capped. We reconstructed bacterial genomes from the metagenomes, and mapped metatranscriptomes to these genomes to determine the activity of oil-responsive functional pathways across a hydrocarbon concentration gradient left by the spill. Studied communities included those from seven highly polluted ‘near-well’ sites around MC252 (0.3 to 2.7 km away), and six ‘distal’ sites that were distributed along a linear transect (10.1 to 59.5 km away)^10^, which followed the prevailing southwesterly deep plume path^6^ (Fig. S1). Distal-most sites were either un-impacted or minimally impacted, based on greatly diminishing plume hydrocarbon concentrations beyond ~30 km^7,20^ and oil-proxy sediment hopane concentrations beyond 40 km from the wellhead^7^. Results provide insights into the genomic potential and in situ transcriptional activity of dozens of spill-responsive bacteria, including more than 20 candidate hydrocarbon degraders.

## Results and Discussion

To establish a site-specific genomic database for metatranscriptome mapping, metagenomic sequences were co-assembled from three genomically representative (Fig. 1) and comparatively well-assembling sediment samples (Table S1) collected 0.5, 0.7 and 0.9 km from MC252. Contigs were binned into genomes aided by compositional information^21^ and differential coverage^22^, which was obtained by mapping metagenomic reads from all 13 sites to the co-assembly. This also enabled us to determine the abundance of co-assembled genomes at each site. Most sequences (66% or 119 Mbp) were classified into 51 bins comprising 57 genomes (Table S2, Dataset S1) associated with the Gammaproteobacteria, Alphaproteobacteria, Deltaproteobacteria and Bacteroidetes (Table S2). A further 4.4% of contigs contained virus-like genes that were mainly associated with the Gammaproteobacteria. The relative abundance of the bacterial genome bins ranged from 0.6 to 13.1% (1.6% on average), and consisted of partial to near complete genomes estimated to be 7 to 97% complete (50% on average; Table S2). Five bins contained 2 to 3 very closely related genomes with partial (insufficient) coverage separation (Fig. S2). Metatranscriptomic sequences, derived from 10 of the same samples as the metagenomes, were then mapped to the co-assembly, generating genome-specific expression profiles up to 33.9 km from MC252 (Dataset S1). This also enabled mRNA hits to be normalized to genome abundance per site to determine gene up/down regulation.

**Figure 1.**
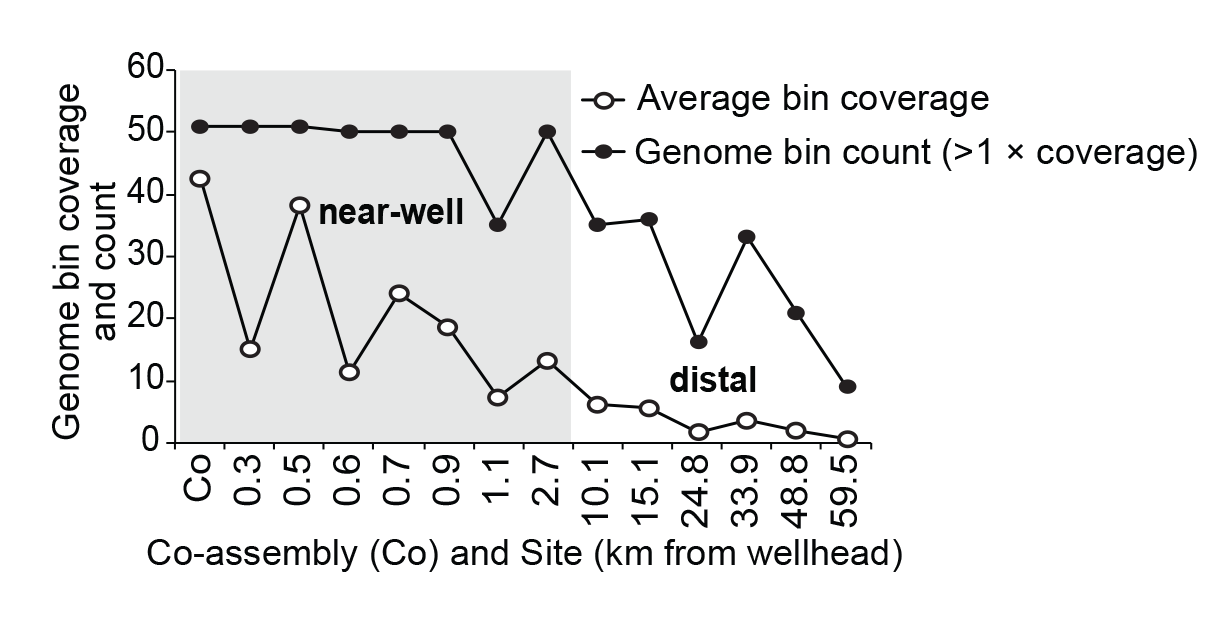
Average genome bin coverage and count (for bins with >1 × coverage) in the co-assembly, and for the same collection of genome bins at each individual site; both decrease noticeably with increasing distance from the wellhead, although some of the genomes were still detectable 60 km away. Distance along the x-axis is not shown to scale.

Small subunit (SSU) rRNA genes and rRNA sequences from the metagenomes and metatranscriptomes were reconstructed using EMIRGE^23^, and co-clustered into operational taxonomic units (OTUs), including rRNA sequences from an additional unpaired metatranscriptome sample collected 1.3 km from MC252. Comparison of the relative abundances of bacterial, archaeal and eukaryotic SSU rRNA gene sequences demonstrated that bacteria near MC252 increased appreciably relative to archaea and eukaryotes (Fig. S3). Gammaproteobacteria predominated near MC252^10^ and exhibited increased species richness (Fig. S3), analogous to the predominance of Gammaproteobacteria observed in the deep-sea plume^14^. While rRNA is not necessarily a good indicator of microbial growth^24^, we found rRNA gene and rRNA relative abundances were generally well correlated for prokaryotes (but not eukaryotes), particularly for bacteria enriched near the wellhead, and for a distally abundant OTU belonging to the Marine Group I Archaea (OTU-15; Fig. 2). These data, along with mRNA expression profiles (Dataset S1), imply spill-responsive bacterial communities were still viable and active ≥75 days after well closure.

**Figure 2.**
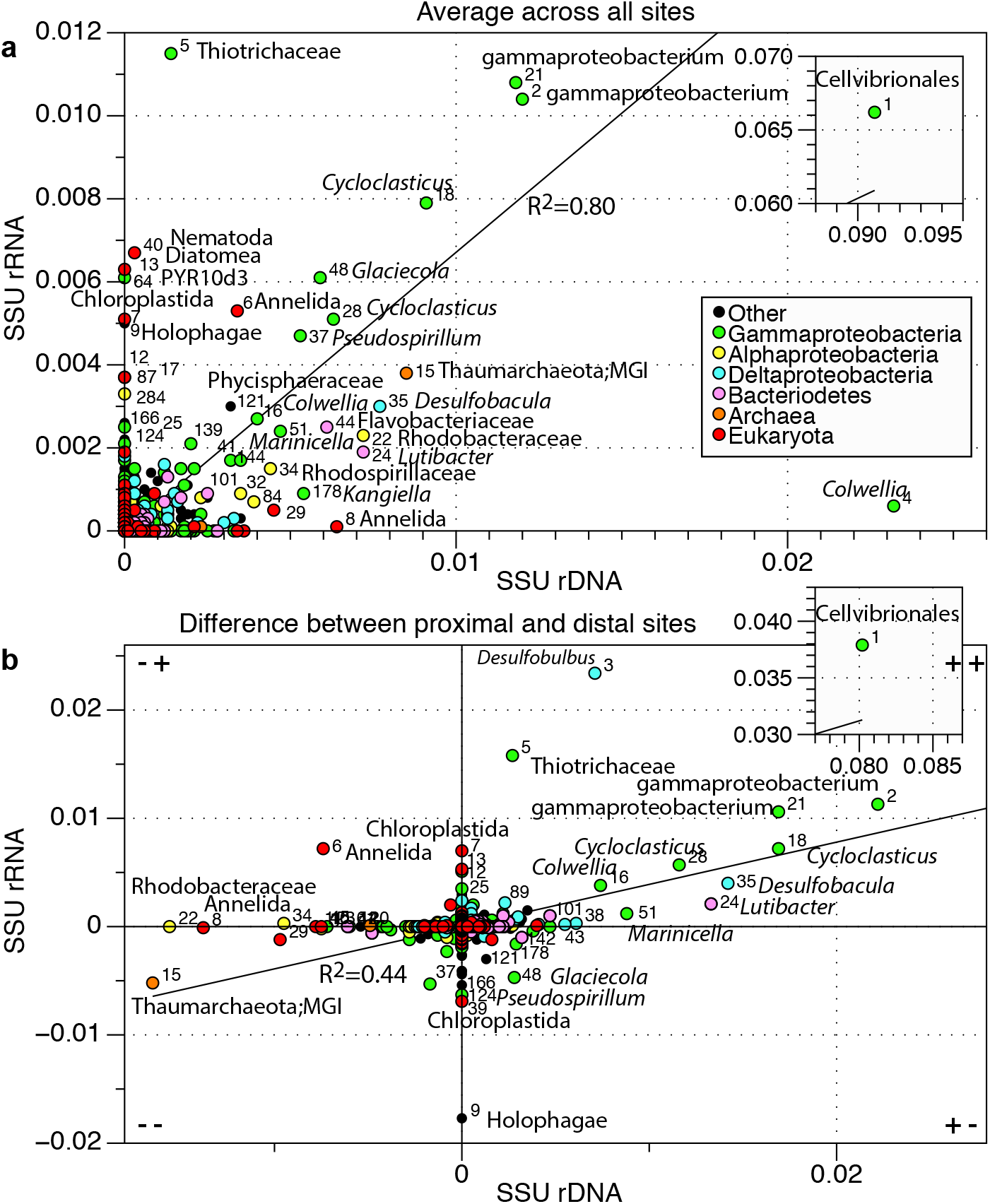
(A) The correlation of EMIRGE 16S rRNA gene and rRNA OTU abundances. (B) The difference between proximal and distal sites. Read clockwise: higher rRNA gene and rRNA in proximal locations (++); higher rRNA gene but lower rRNA proximally (+-); lower rRNA gene and rRNA proximally (--); and lower rRNA gene but higher rRNA proximally (-+). (A-B) OTU numbers are given besides taxa points. Insets show the highly abundant OTU 1. Based on abundance and phylogenetic affiliation, OTU 1 corresponds to the *Ca.* Cellvibrionales GSC11-15 genomes (93% identity to *P. hydrocarbonoclasticus*). It also shares 100% identity (ID) with an iTag sequence (>97% similar to Greengenes OTU 248394) from a highly abundant uncultured gammaproteobacterium previously identified in the contaminated near-well sediments 10. Thiotrichaceae OTU 5 corresponds to *Ca.* Thiotrichaceae GSC1 (99% ID to *Ca.* Halobeggiatoa sp. HMW-S2528); *Cycloclasticus* OTUs 18 and 28 respectively correspond to *Ca.* Cycloclasticus GSC8 and GSC9–10 (respectively 98% and 95% ID to *Cycloclasticus zancles* 78-ME); *Colwellia* OTU 4 corresponds to *Ca.* Colwellia GSC4 and GSC9 (99% ID to *Colwellia psychrerythraea* 34H), and *Colwellia* OTU 16 corresponds to *Ca.* Colwellia GSC5-6 (97% ID to *Colwellia* sp. MT41).

Likewise, bacterial genome bins were highly and exponentially enriched near MC252 (Fig. S4, Table S3); the average genome read-coverage was only 0.7 to 6.2× across widespread sites 10-60 km away, but was 7.3× to 42.6× among sites within 2.7 km of MC252 (Fig. 1, Fig. S5). The genomes were found across all near-well sites, as well as at many less impacted distal locations, depicting a remarkably uniform community response across vast areas of the seafloor. Importantly, these oil responsive bacteria were universally distributed, with detection of 54 to 100% of contigs from each of the 51 bins across all 13 sites (Table S4), with 6 Gammaproteobacteria and 2 Bacteroidetes bins present at sites spanning the entire 60 km (Table S5, Fig. S6). These data suggest that, in addition to a few highly abundant OTUs previously identified in the plume and along the seafloor^10,15,18^, there was also a widespread community-level response. Bacteria at the furthermost outreaches of our study area were either responding to low levels of hydrocarbon contamination or were background community members.

Many of the DWH seafloor genomes were phylogenetically clustered (52%, co–binning excluded). Clusters in each of our sampled proteobacterial classes and the Bacteroidetes shared average amino acid identities of 60-86%, which broadly equates to genus or family level relatedness^25^. Of these, the Gammaproteobacteria exhibited 5 distinct inter-bin clusters (Table S6). Prevalent phylogenetic clustering suggests strong habitat selection for traits shared among genetically similar organisms^26^. Gammaproteobacterial clusters included genomes related to sulfur-oxidizing *Candidatus* Halobeggiatoa^27^ (Fig. 2), and to hydrocarbonoclastic *Colwellia*, *Cycloclasticus* and *Porticoccus* (Cellvibrionales) species^28-30^ (Table S7, Fig. S7). Of these, *Colwellia* and *Cycloclasticus* were typical plume genera^14^. When compared with cultivated representatives, EMIRGE-reconstructed 16S rRNA gene sequences related to *Cycloclasticus* and *Porticoccus* belonged to distinct DWH seafloor clades (Fig. S8), while extremely diverse sequences were associated with the *Colwellia*-*Thalassomonas*-*Glaciecola* group (Fig. S8).

The phylogenetic characteristics of the proximal sediment communities suggested a strong potential for sulfur and hydrocarbon metabolism; also supported by gene content and gene expression profiles. Among the most highly expressed genes were those exhibiting significant differential expression between proximal and distal sites for sulfur, hydrocarbon and nitrogen metabolism (Fig. 3). An increase in genes involved in N metabolism was previously identified in these oil-polluted sediments^10,31^. Our data show these genes – involved in the denitrification pathway and nitrite/hydroxylamine oxidation/reduction – were significantly up-regulated at proximal sites. While denitrification-related activity was associated with several Gammaproteobacteria, it was mostly linked to anaerobic sulfur oxidation by two Halobeggiatoa-like Thiotrichaceae (GSC1 and 3; Fig. 3 and S9). These Thiotrichaceae also expressed of hydroxylamine reductase genes (EC 1.7.2.6), which they likely used as a supplemental method for reducing nitrite^32^ to ammonia, although a hydroxylamine reductase required for the final step (hydroxylamine reduction to ammonia) was not detected in any Thiotrichaceae genome bin. Concomitant up-regulation of oxidative phosphorylation genes reflects the classic oscillating aerobic-anaerobic lifestyle of this group of bacteria^33^. Genes involved in Thiotrichaceae sulfur oxidation, and deltaproteobacterial sulfate reduction, were both significantly up-regulated near MC252, which implies an active spill-promoted sulfur cycle formed in the top 1 cm of seafloor sediment.

**Figure 3.**
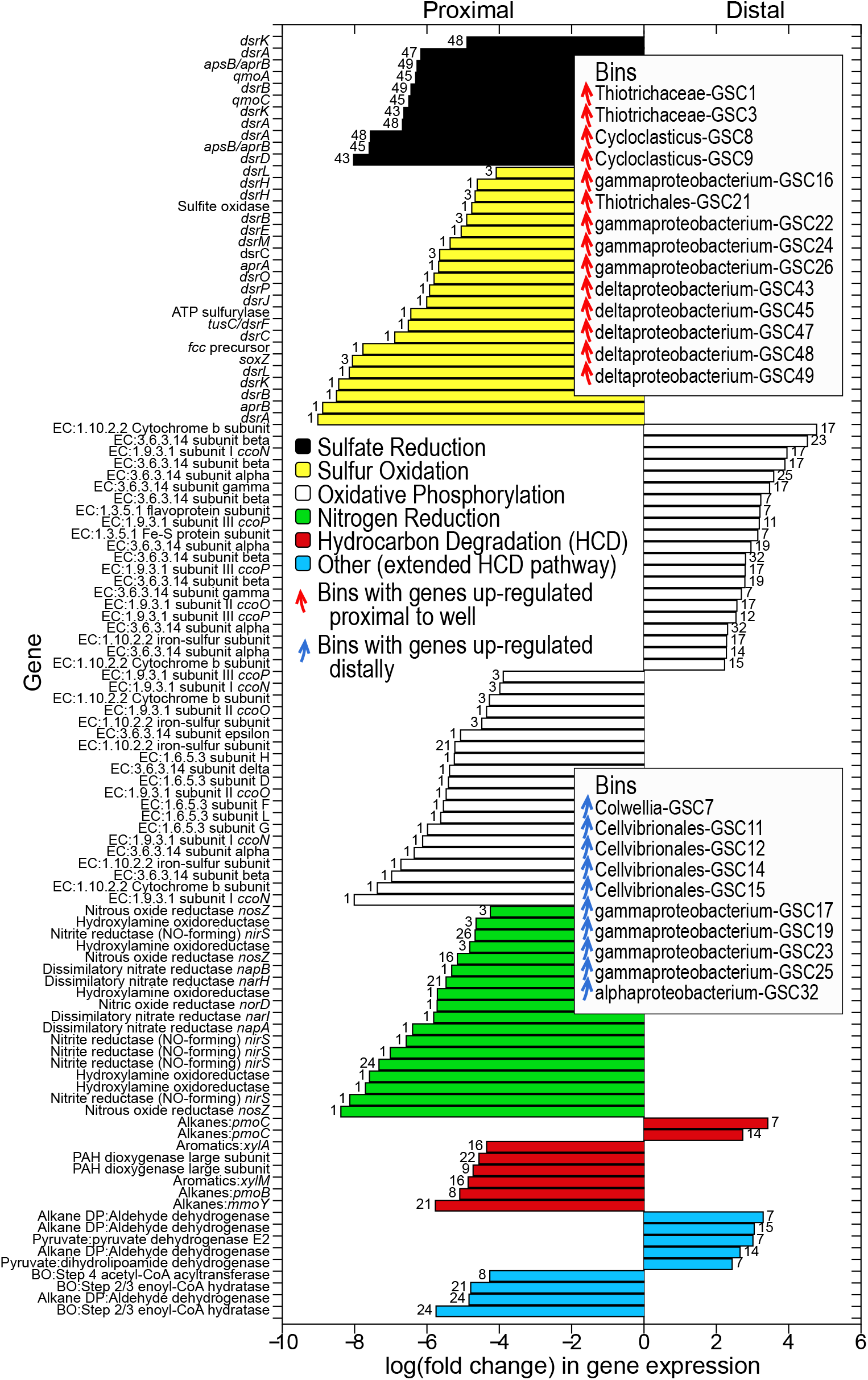
Differentially expressed genes at proximal and distal sites associated with hydrocarbon degradation, oxidative phosphorylation, and S cycling and N reduction pathways. Genome bin numbers (GSC) are given beside each bar. Abbreviations: degradation pathway (DP); beta oxidation (BO).

Genes indicative of hydrocarbon degradation (HCD) were concentrated in the Gammaproteobacteria (179 genes or 66%; Table 1 and Table S8). Half of the bacterial genomes (n=25) contained genes associated with the degradation of hydrocarbons, most belonging to aerobic pathways (Table S8). These genes were observed almost exclusively in additive combinations targeting: (1) *n*-alkanes, (2) *n*-alkanes + aromatics, or (3) *n*-alkanes + aromatics + PAHs (Figs S4 and S9), whereby alkane degradation potential is the common denominator. As such, genomes with genes for *n*-alkane degradation were cumulatively the most abundant near MC252 (Fig. S4). We observed the expression of several genes involved in the aerobic degradation of these 3 hydrocarbon classes across multiple seafloor sites, although a greater proportion of genes targeting *n-*alkane versus (poly)aromatic substrates were expressed at distal sites (Fig. 4). Overall a greater number of genes associated with hydrocarbon degradation were up-regulated proximally (Fig. 3), likely due to the higher average concentration of hydrocarbons near MC252 (Fig. S4)^10^.

**Table 1.**
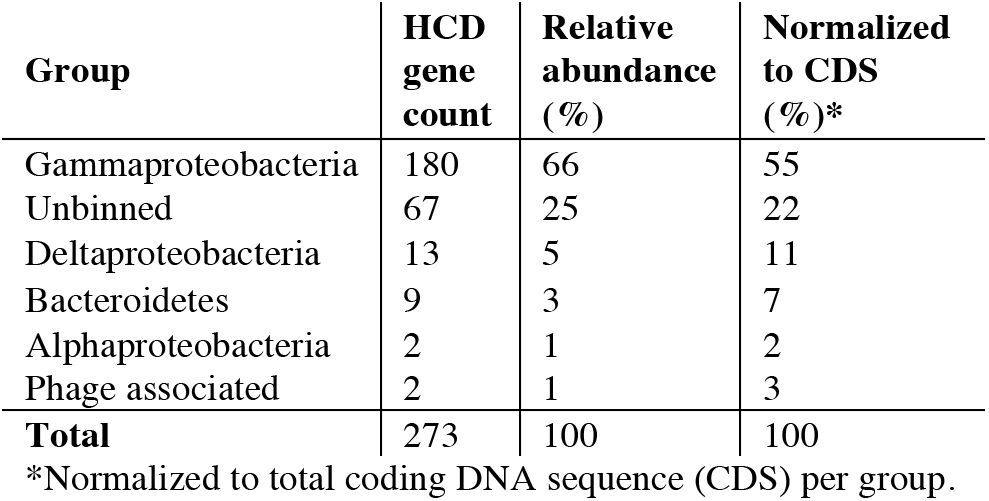
Hydrocarbon degradation (HCD) gene distributions.

| **Group** | **HCD**<br>**gene**<br>**count** | **Relative**<br>**abundance**<br>**(%)** | **Normalized**<br>**to CDS**<br>**(%)\*** |
| --- | --- | --- | --- |
| Gamma proteobacteria | 180 | 66 | 55 |
| Unbinned | 67 | 25 | 22 |
| Delta proteobacteria | 13 | 5 | 11 |
| Bacteroidetes | 9 | 3 | 7 |
| Alpha proteobacteria | 2 | 1 | 2 |
| Phage associated | 2 | 1 | 3 |
| **Total** | 273 | 100 | 100 |

\*Normalized to total coding DNA sequence (CDS) per group.

**Figure 4.**
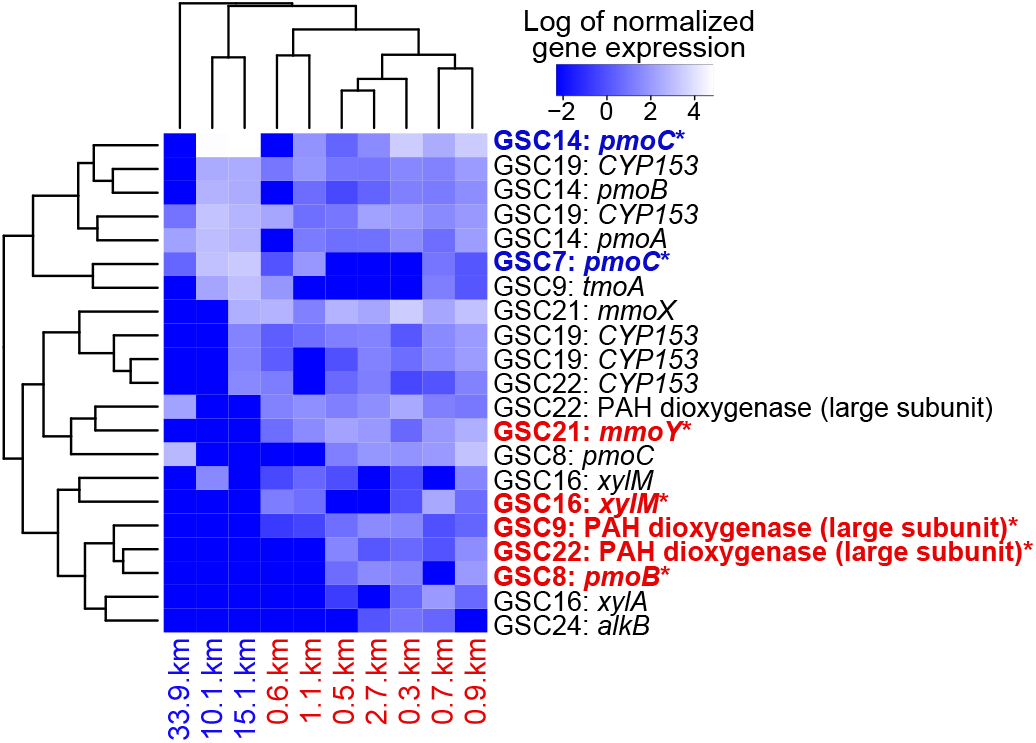
Spatial expression profiles of genes associated with hydrocarbon degradation. Genome bin and gene identities are given for each row. *Significantly differentially expressed genes at proximal (red font) versus distal (blue font) sites.

The genes that were differentially expressed for aerobic HCD and associated oxidative phosphorylation belonged to 2 largely distinct groups of Gammaproteobacteria (Fig. 3). The first group exhibited higher expression of genes at proximal sites involved in the degradation of short-to-mid chain length *n*-alkanes, aromatics and PAHs (*Ca.* Cycloclasticus GSC8 and GSC9, *Ca.* Thiotrichales GSC21, gammaproteobacterium GSC16, and Chromatiales/Thiotrichales-relative GSC22). The second up-regulated genes at distal sites associated with short-to-mid chain *n*-alkanes (*Ca.* Colwellia GSC7, and *Ca.* Cellvibrionales GSC14). While both groups of bacteria were active across wide distances, the second group appeared to be more active at 10 and 15 km distances (Fig. 4) where total petroleum hydrocarbon and *n*-alkane concentrations were generally lower^10^ (Fig. S4).

We identified 3 candidate PAH degraders (*Ca.* Cycloclasticus GSC9, Chromatiales/Thiotrichales-related GSC22, and *Ca.* Cellvibrionales GSC15), of which GSC9 and GSC22 exhibited active expression of PAH dioxygenases, particularly near-well. Related *Cycloclasticus* species and *Porticoccus hydrocarbonoclasticus* (Cellvibrionales) are known for their ability to degrade various PAHs^28,29^. All 3 candidate PAH degraders had broadly equivalent spatial abundances (Fig. S5), and each genome has 17-24 subunits (large and small) of diverse ring-hydroxylating dioxygenases that, except for 2, closely resemble dioxygenases used for the oxidation of PAHs and other aromatic hydrocarbons (Fig. S9 and Dataset S1)^34^. As previously observed in *Cycloclasticus* genomes^35,36^, each of our 3 DWH genomes had a greater proportion of large over small PAH dioxygenase subunits (58-63% in genomes and 64% unbinned).

PAH dioxygenase sequences from the GSC9, GSC22 and GSC15 genomes chiefly resembled naphthalene dioxygenases (23 large and 17 small subunits), while the remainder (20 large and 11 small subunits) were more closely related to biphenyl/benzene, anthranilate/ (ortho-halo)benzoate or pyrene dioxygenases (Fig. S10 and Table S8). PAH dioxygenases produce dihydrodiols^37^. Canonical naphthalene, anthranilate and pyrene *cis*-dihydrodiol dehydrogenases (EC 1.3.1.29, EC 1.3.1.49) were not evident in our DWH genomes. However, all 3 genomes possess *cis*-2,3-dihydrobiphenyl-2,3-diol dehydrogenase (*bphB*) and 2,3-dihydroxybiphenyl 1,2-dioxygenase (*bphC*) genes. BphB (EC 1.3.1.56) is a multi-substrate enzyme that acts on biphenyl-2,3-diol and a wide range of PAH dihydrodiols^37^, including those relevant to MC252 oil, such as naphthalene 1,2-dihydrodiol, and phenanthrene and chrysene 3,4-dihydrodiols^8^. BphC degrades the catechol product of BphB. These bacteria appear to use a universal pathway for the catabolism of early PAH degradation products, as opposed to distinct dihydrodiol dehydrogenases per PAH substrate identified in a collection of Gulf of Mexico seawater-associated Gammaproteobacteria^38^.

Genetic mechanisms for BTEX degradation were previously found to be enriched at highly contaminated seafloor sites^10^. Through assembly and bin assignment, we were able to link genes used for toluene, xylene and benzene degradation with at least 4 Gammaproteobacteria: GSC9, GSC22, GSC16 (related to *n*-alkane-degrading gammaproteobacterium HdN1^39^) and GSC24 (related to Oceanospirillales *Hahella chejuensis*). Collectively these 4 genomic bins represented 8% of the average genome abundance at the proximal sites (Fig. S4). All 4 bins had genes encoding xylene monooxygenase-like enzymes (Xyl), which oxidize toluene and xylenes^40^. *xylAMM* genes were expressed by GSC16 and was significantly higher at the proximal sites (Fig. 3). In contrast, GSC9 demonstrated greater expression, albeit not significantly, of a gene (*tmoA*) encoding part of a largely unbinned aerobic toluene-4-monoxygenase system at distal sites, suggesting toluene/xylene metabolism very likely occurred across the seafloor, although the organisms and mechanisms varied. GSC9 and GSC22 also had genes encoding multicomponent phenol hydroxylase like enzymes (Dmp), which oxidize phenol, benzene and toluene^41^. Further to these mechanisms, the 3 candidate PAH degraders (GSC9, GSC22 and GSC15) had putative benzene 1,2-dioxygenases (also similar to biphenyl dioxygenases) and catechol 2,3-dioxygenases (EC 1.13.11.2), suggesting these taxa could generate catechol by benzene oxidation, which could then be converted into 2-hydroxymuconate-semialdehyde, and sequentially transformed into pyruvate (Fig. S9). While hydrocarbon degradation mechanisms identified in these surface sediment communities were overwhelmingly aerobic, there were a few exceptions in sequence data not binned to genomes. These include a single set of anaerobic ethylbenzene dehydrogenase genes (*ebdACBA*); and benzylsuccinate synthase genes (*bssCAB* and *bssCA*), which can be used for anaerobic toluene oxidation^42^, and were observed in the lower anaerobic layer of seafloor sediments polluted with oil from MC252^43^.

Widespread evidence for alkane oxidation was associated with 3 different mechanisms that target gaseous C2-C4 short-chain and liquid C5-C10 mid-chain alkanes. Genes associated with the oxidation of both short to mid length *n*-alkane groups were expressed across at least 34 km of the Gulf of Mexico seafloor (Fig. 4). Of these, Alk and CYP153 enzymes act on mid-chain alkanes from pentane to decane (C5-C10)^44^. Transmembrane 1-alkane monooxygenase (AlkB ± AlkGT rubredoxin/rubredoxin reductase) and membrane-bound cytochrome P450 CYP153 (± ferredoxin/ferredoxin reductase) hydroxylases genes were both present, although CYP153 were more commonly expressed. Pathway analysis of 14 key gammaproteobacterial bins with alkane hydroxylases suggests 1-alcohol generated by alkane oxidation could be converted sequentially to aldehydes by alcohol dehydrogenase (EC 1.1.1.1), carboxylates by aldehyde dehydrogenase (NAD, EC 1.2.1.3), and acetyl-CoA via beta-oxidation (Fig. 3).

Also present were genes resembling particulate and soluble methane or ammonia monooxygenases. These constitute a group of related multi-substrate enzymes that preferentially target methane or ammonia^45^. They can be used by methanotrophs, in the absence of methane, to oxidize short *n*-alkanes, namely C2-C4 gases (ethane, butane, propane) and C5 liquid (pentane)^46,47^. Longer C6-C8 alkanes, can also be used, albeit at an appreciably slower rate^46^. In comparison, *Mycobacterium* strains preferentially use soluble monooxygenases to oxidize alkanes^48^. We recovered a diverse group of 5 genomes with particulate methane monooxygenase like genes (*Candidates* Colwellia GSC7, Cycloclasticus GSC8, Cellvibrionales GSC14 and Thiotrichales GSC21, and gammaproteobacterium IMCC2047 relative GSC18). Thiotrichales GSC21 also had genes encoding the components of a soluble monooxygenase (sMMO), which is used in place of the particulate enzyme (pMMO) under copper limiting conditions^49^. All 5 genomes lack evidence for methanol or hydroxylamine oxidation, suggesting an inability to utilize the products of methane or ammonia oxidation. However, all had the genetic capacity to convert 1-alcohols generated from *n*-alkane oxidation to acetyl-CoA (Fig. S9). We therefore predict that these bacteria primarily used pMMO ± sMMO to oxidize *n-*alkanes. The lack of dedicated methanotrophs plausibly reflects the absence of trapped methane in the post-spill sediments, and is consistent with the absence of methane in the late-stage plume^50^.

We found that both types of methane monooxygenase like genes were expressed across multiple seafloor locations; three bacteria expressed particulate *pmo* genes (GSC7, GSC8, GSC14), while Thiotrichales GSC21 expressed soluble *mmo* genes (Fig. 4). GSC21 co-expressed a short chain alcohol dehydrogenase gene, resembling a NADH-dependent butanol dehydrogenase gene (*bdhA*), which it may have used to metabolize 1-alcohol produced by alkane degradation (Fig. S9). We detected expression of parts of the downstream alkane degradation pathway by GSC7 and GSC8 (Dataset S1). All genes comprising the pathway from 1-alcohol to the first steps in the beta-oxidation pathway were expressed by GSC14, including multiple NAD-dependent alcohol dehydrogenase (EC 1.1.1.1) and aldehyde dehydrogenase genes.

## Conclusions

Gulf of Mexico microorganisms are naturally exposed to oil seeps^5,13^ and frequent spills^2^. Our genomic and metabolic reconstruction of oil-impacted communities distributed across the seafloor indicates that a large common collection of bacteria responded to the DWH spill, many of which possessed hydrocarbonoclastic potential. A large degree of apparent functional redundancy among HCD strategies suggests that the Gulf of Mexico harbors functionally robust communities that are well poised to take advantage of petroleum hydrocarbon influxes. We observed a strong environmental selection preference for genetically similar organisms, implying that the preservation or sharing of opportunistic hydrocarbonoclastic (and S-oxidizing) traits was important among these DWH organisms. Due to the substrate promiscuity of many hydrocarbon-degrading enzymes^34,41,44,48^, it is unclear whether actively transcribed genes resulted in competition for the same substrates or niche differentiation. Nevertheless, our results show that several closely related hydrocarbon-degrading genes were concomitantly expressed, and that individual bacterial populations appeared to occupy more than one niche by co-utilizing functionally distinct hydrocarbon-degrading genes.

## Methods

### Sampling and nucleic acid sequencing

Thirteen seafloor sediment cores were collected between 28 September and 19 October 2010 at radially distributed locations around the capped MC252 wellhead (x7 cores between 0.3 and 2.7 km from MC252), and along a distal southwesterly linear transect (x6 cores between 10.1 and 59.5 km from MC252) (Fig. S1 and Table S1)^10^. The outer surfaces of 0 to 1 cm deep cores were removed prior to DNA and RNA extractions^10^. DNA extraction and whole genome shotgun (WGS) sequencing are described by Mason et al.^10^. Briefly, DNA extracted from the cores was fragmented and prepared for sequencing using Illumina's TruSeq DNA Sample Prep Kit. Each library was sequenced on a full HiSeq2000 lane at the Institute for Genomics and Systems Biology's Next Generation Sequencing Core (Argonne National Laboratory). This yielded ~18 Gb of sequence per sample with 2 x 101 bp reads and insert sizes of ~135 bp. Low quality reads were trimmed using Sickle v. 1.29 with a quality score threshold of Q=3, or removed if trimmed to <80 bp long (<u>https://github.com/najoshi/sickle</u>).

RNA was extracted in duplicate from cores using 0.5 g of sediment for each replicate. A modified hexadecyltrimethylammonium bromide (CTAB) extraction buffer was used as previously described^51^. Duplicate extracts were pooled, and purification was carried out using the Qiagen AllPrep DNA/RNA Mini Kit with on-column DNase digestion using the RNase-Free DNase Set (Qiagen, Valencia, CA). RNA from samples with low yields (<150 ng RNA: 0.3 km, 1.1 km, 10.1 km, 15.1 km and 33.9 km samples) was amplified using the Ambion MessageAmp II aRNA Amplification Kit (Foster City, CA). RNA was converted into double stranded cDNA using the SuperScript Double-Stranded cDNA Synthesis Kit with random hexamers (Invitrogen, Carlsbad, CA). Double stranded cDNA was prepared for sequencing using the TruSeq Nano DNA kit (Illumina, San Diego, CA). Prepared libraries (~440 bp long) were sequenced using an Illumina HiSeq 2500 at the Yale Center for Genomic Analysis, with 3 libraries per lane, and generating 150 bp paired-end reads. Adapter sequences were removed using Cutadapt^52^, and reads were trimmed with Trimmomatic^53^ (sliding window quality score ≥15) and removed if shorter than 60 bp.

### Metagenome assembly

Metagenomic sequences from each sample were first assembled individually using the IDBA-UD v. 1.1.0 metagenome assembler^54^. Consolidation and improved genome and HCD gene recovery was achieved via a co-assembly of 3 representative samples (0.5, 0.7, 0.9 km from MC252), due to high compositional similarity among near-well communities. IDBA-UD was used for the co-assembly with an optimal kmer range of 45 to 75 and step size of 15. Improved recovery of the highly abundant GSC11 genome was achieved through selective re-assembly from the 0.5 km metagenome using Velvet (kmer size = 63, expected kmer coverage = 147, kmer coverage range = 92 to 225)^55^.

### Genome binning, annotation, completion estimates and comparisons

To bin contigs ≥2 kbp long, we used the multi-parameter approach previously described by Handley et al.^56^. To better separate closely related genomes using emergent self-organizing maps (ESOM), contig tetranucleotide frequencies were augmented with the coverage of that contig in each spatially distinct sample (Fig. S11) based on the approach of Sharon et al.^22^. The differential coverage of contigs was determined by mapping reads to co-assembled contigs using bowtie2^57^. Contig coverages per sample were scaled to 1 for ESOM. Genome bin abundance heat and line plots were created using gplots and ggplot2 packages in R, respectively.

Contigs were annotated using the Integrated Microbial Genomes (IMG) pipeline^58^, and the Rapid Annotations using Subsystems Technology (RAST) Server^59^. Predicted protein sequences of potential HCD genes were searched against the National Center for Biotechnology Information's (NCBI's) Conserved Domain Database^60^.

Genome completion was estimated based on the presence of 107 single copy core genes^61^, excluding *glyS*, *proS*, *pheT* and *rpoC*, which were missing or poorly recovered in this study and according to Albertson et al.^62^. Core genes were detected using HMMER3 with the default cutoff^63^. Estimates were similar to those obtained using AMPHORA2 with a set of 31 universal bacterial protein-coding housekeeping genes^64^.

Genomes were compared using average amino acid identities (AAIs) including only BLASTp matches that shared ≥30% identity over an alignable region of ≥70% sequence length. 16S rRNA gene sequence and RecA and PAH dioxygenase predicted protein sequences were compared using MEGA6^65^ ClustalW alignments and neighbor-joining or maximum-likelihood trees.

### Transcriptome read mapping

Between 21 and 58 million trimmed paired-end reads were mapped to the co-assembly using Bowtie2^57^ with --end-to-end and --sensitive settings. Alignments were sorted with Samtools^66^. Hits were enumerated and filtered using htseq-count in HTSeq v. 0.6.1p1^67^ with default settings. Counts were normalized to gene length and reads per sample by a modification of the approach described by Mortazavi et al.^68^, whereby normalization was to Reads Per Kilobase per Average library size (RPKA; average = 44 million). To determine whether gene expression was up or down regulated spatially, RPKA values were also normalized to the genome coverage in each sample (RPKAC, an we required that at least one sample per gene had at least 10 mapped reads (un-normalized). To compare differential gene expression between proximal and distal sites we used edgeR^69^ on un-normalized data with RPKAC values supplied as an offset matrix of correction factors to the generalized linear model. Sample sizes were supplied as total library sizes.

### Ribosomal RNA sequence assembly and clustering

To investigate beta diversity near full-length small subunit (SSU) rRNA (gene) sequences were reconstructed using the reference-guided Expectation Maximization Iterative Reconstruction of Genes from the Environment (EMIRGE) method^23^. WGS samples were rarefied to an equal depth of 64 million paired-end reads. Transcriptome samples were rarefied to between 22 and 32 million reads after first pre-selecting SSU rRNA specific transcriptome kmers using bbduk (<u>http://sourceforge.net/projects/bbmap</u>) and the SILVA SSU rRNA database^70^. Sequences were reconstructed over 80 iterations using EMIRGE with the SILVA 111 SSU rRNA database. Gapped and chimeric sequences were removed using 64-bit USEARCH v. 8.0 with the RDP Gold v. 9 database (http://drive5.com/). RNA and RNA gene sequences were co-clustered at ≥97% similarity into operational taxonomic units (OTUs) using -cluster_otus (32-bit USEARCH v. 8.1)^71^ after sorting by size. OTU representative sequences were identified by 64-bit USEARCH global alignment to the full SILVA 111 SSU rRNA database, and by RDP classifier v. 2.6^72^ with a 0.8 bootstrap cutoff.

### Sequence accession

Metagenomic and EMIRGE assemblies, and transcriptome reads are accessible via NCBI BioProject's PRJNA258478 and PRJNA342256, respectively. Metagenome annotations are accessible via IMG ID 3300003691.

## Acknowledgements

This work was supported by Alfred P. Sloan Foundation and Exxon-Mobile grants awarded to JAG, and a Royal Society of NZ Rutherford Discovery Fellowship awarded to KMH. Partial support was provided by the U.S. Dept. of Energy under contracts DE-AC02-06CH11357 (ANL) and DE-AC05-76RL01830 (PNNL). We thank Christian Sieber (JGI) for transcriptome rRNA sequence assembly, and acknowledge resources provided by the University of Chicago Research Computing Center, NERSC, and the University of Auckland NeSI high-performance computing facilities and Centre for eResearch.

## Supplementary figure, table and dataset legends

Figure S1. (a) Location of well MC252, and (b) sampling locations. Samples >3 km from MC252 are shown in blue, while near-well samples are shown in red (inset).

Figure S2. Coverage and GC content of contigs within each bin. Y-axes have different scales (LHS of each plot). *Compositionally similar co-binned genomes with imprecise coverage separation.

Figure S3. Phylogeny based on rRNA genes reconstructed from rarefied data, (a) Phylogeny per site. Eukaryota are relatively abundant in distal sites (23-52% versus 6-34% near-well). Bacteria, particularly Gammaproteobacteria increase in abundance relative to both Eukaryota and Archaea. (b) Stacked bar chart of Gammaproteobacteria rRNA gene sequence coverage per site based on RDP genus level designation or higher. The average number of unique designations is 13 ± 2 (1 standard deviation) for near-well sites, twice that for distal sites (7 ± 3). (c) Neighbor-joining tree depicting phylogenetic 16S rRNA gene diversity among the Gammaproteobacteria for near-well (red) and distal samples (blue), compared with reference sequences (black). Genus (italics) and order (bold) names are given for reference sequence clusters. Gammaproteobacterial richness at near-well sites was 317 (15,000x total 16S rRNA gene coverage), and the richness at distal sites was 152 (6,000x total 16S rRNA gene coverage).

Figure S4. (a) Boxplots of petroleum hydrocarbon and carbon concentrations in samples 0-3 km (near-well) and 10-60 km (distal) from the MC252 well-head. Abbreviations: (total) petroleum hydrocarbons, (T)PHC; weight, wt. *Concentrations enriched near-well. (b) Summed per site abundance of genome bins possessing genes associated with hydrocarbon degradation. Points are fitted with exponential curves. Bin numbers are listed in boxes central to each plot. Points in yellow represent the combined abundance of bin numbers given in black and yellow.

Figure S5. Average genome bin coverages per site determined by mapping reads to the co-assembly. Error bars = 1 standard deviation. Samples contributing to the co-assembly (white points) are bolded and points shown in yellow. Plots are organized by phylogenetic group: Gammaproteobacteria (green), Alphaproteobacteria (yellow), Deltaproteobacteria (blue), Cytophaga-Flavobacterium-Bacteroides group (CFB, pink),
and unknown Bacteria (grey).

Figure S6. Heatmap and hierarchical clustering of genome bin abundance per site or Co-assembly (Co). Samples contributing to the co-assembly are bolded. Abundances ≥10 are assigned the same color (dark red). Bins (RHS) are colored by taxonomic group: Gammaproteobacteria (green), Alphaproteobacteria (yellow), Deltaproteobacteria (blue), Cytophaga-Flavobacterium-Bacteroides group (CFB, pink), and unknown Bacteria (grey).

Figure S7. Maximum-Likelihood tree showing the phylogenetic relatedness among recombinase A (RecA) predicted protein sequences from our seafloor genome bins (green) and reference organisms (black). Reference sequence GenBank accession numbers are in parentheses. Tree construction employed 190 amino acid positions and 500 bootstrap replicates.

Figure S8. Maximum-Likelihood trees showing the phylogenetic relatedness of rRNA genes sequences from near-well (red) and distal (blue) sites that are similar to (a) *Cycloclasticus*, (b) *Porticoccus*, and (c) *Colwellia* species. Sequence identifiers are in parentheses. Norm Priors (NP) denote abundances per sample. Tree construction employed near full-length 16S rRNA genes and 500 bootstrap replicates. 16S sequences related to these hydrocarbonoclastic genera were recovered from rarified sequence data up to ~10 km from MC252 for *Cycloclasticus* and *Porticoccus*, or 33.9 km in the case of *Colwellia*.

Figure S9. (a) Idealized cell-schematic showing genes GSC1 likely uses to encode for denitrification and sulfide oxidation. Genes present are in grey, while those absent are in red. (b) Cell schematic of 14 gammaproteobacterial genome bins with the metabolic potential to degrade hydrocarbons. Genes present are in black; those missing are in red. Subunits are listed in order present in genomes, and separated by a dash if on different contigs. Substrates (and their products) are categorized by color: C2-C10 alkanes (dark blue), methane (light blue), aromatic hydrocarbons (orange), PAHs (purple), nitrogen species (green), and other (black). All genome bins contain (near) complete beta oxidation pathways that can be used to oxidize hexadecanoate or fatty acyl-CoA esters intermediates to acetyl-CoA.

Figure S10. Maximum-Likelihood tree depicting the phylogenetic relatedness among predicted protein sequences of (near) full length candidate PAH dioxygenases alpha subunits from our genome bins (green) compared with reference naphthalene, pyrene and anthranilate PAH dioxygenases and biphenyl, benzene and (ortho-halo-)benzoate aromatic dioxygenases. GenBank accession numbers for reference sequences are in parentheses. Tree construction employed 340 amino acid positions and 500 bootstrap replicates.

Figure S11. (a) ESOM of co-assembly constructed using tetranucleotide frequencies of 5 kbp long genomic fragments and differential coverage. Bin numbers are shown in red. Dark lines demark cluster edges. Collections of small clusters are contigs containing virus-associated genes. (b-e) ESOM with data points representing 2 kbp long genomic fragments and colored by bin (see key).

Table S1. Sequence data input (trimmed), and IDBA-UD assembly summary.

Table S2. Genomic bin characteristics.

Table S3. Pearson's correlations between distance and (hydro)carbon concentrations or genome coverage.

Table S4. Percentage with mapped reads at each site.

Table S5. Average mapped-read genome bin coverages per site.

Table S6. Pairwise average amino acid identities (AAI) shared between genome bins.

Table S7. Average amino acid identities (AAI) shared between genome bins (left) and reference genomes (top).

Table S8. Summary of candidate hydrocarbon degradation genes distributed among 25 bacterial genome bins, GSC52 and the unbinned fraction.

Dataset S1. Contigs per bin; key genes involved in hydrocarbon degradation, nitrate reduction and sulfur oxidation; and mRNA read counts for key genes.

